# Predicting the Specificity-Determining Positions of Paralogous Complexes

**DOI:** 10.1101/2021.01.26.428202

**Authors:** Tülay Karakulak, Ahmet Sureyya Rifaioglu, João P.G.L.M. Rodrigues, Ezgi Karaca

## Abstract

Due to its clinical relevance, modulation of functionally relevant amino acids in protein-protein complexes has attracted a great deal of attention. To this end, many approaches have been proposed to predict partner-selecting, i.e., specificity-determining positions in evolutionarily close complexes. These approaches can be grouped into sequence-based machine learning and structure-based energy-driven methods. In this work, we assessed these methods’ ability to map the specificity-determining positions of Axl, a receptor tyrosine kinase involved in cancer progression and immune-related diseases. For this, we used three sequence-based predictors – SDPred, Multi-RELIEF, and Sequence Harmony – and a structure-based approach by utilizing HADDOCK and extensive molecular dynamics simulations. As a result, we show that (i) sequence-based methods overpredict the number of specificity-determining positions for Axl complexes and that (ii) combining sequence-based approaches with HADDOCK provides the most coherent set of predictions. Our work lays out a critical study on the comparative performance specificity-determining position predictors. It also presents a combined sequence-structure-based approach, which can guide the development of therapeutic molecules capable of combatting Axl misregulation in different types of diseases.

## INTRODUCTION

Functional identification of proteins is integral to understand the grounds of innate cellular processes. Several computational tools have been deployed to annotate protein function from ever-accumulating protein sequences (Friedberg 2006). These approaches aim to define functionally important domains or residues through comparative sequence analysis (Whisstock & Lesk 2004). Resolving functionally key amino acids is particularly interesting, as modulation of these residues holds a great potential to design novel protein-based therapeutics (Moll et al. 2016). Such key residues can be identified by searching for conserved positions across different species. Alternatively, given a single species, one could look for the differential amino acid positions within a family of very closely-related proteins, i.e., paralogs (Gogarten & Olendzenski 1999; Mirny & Gelfand 2002; Chagoyen et al. 2016). In such a family, the differentially mutated residues could have evolved to regulate protein interaction networks and act as specificity-determining positions (SDPs) (Rausell et al. 2010; Sloutsky & Naegle 2016). The fibroblast growth factor (FGF) and their receptors (FGFRs) are an example of paralog protein families, where small amino acid changes determine the interaction specificity. FGFs mediate a diverse range of developmental processes through their specific interactions with different FGFRs (Beenken & Mohammadi 2009). The human genome encodes 18 FGFs, leading to 126 possible different FGF-FGFR interactions (Kuriyan et al. 2012). However, only a few of these complexes are productive, establishing the basis of selective FGF-mediated signaling (Beenken & Mohammadi 2009). As a result, accurate predictions of SDPs in the FGF family would help resolve and control such selective interactions.

### Sequence-based SDP predictions

Several sequence-based SDP prediction methods have been proposed in the last three decades (Pirovano et al. 2006; Chakrabarti & Panchenko 2008; Chakraborty & Chakrabarti 2015; Chagoyen et al. 2016). These methods rely on the application of different machine learning techniques, which can be grouped into entropy-, evolution-, and feature-based (Teppa et al. 2012). The majority of these methods expand on the use of a precalculated multiple sequence alignment (MSA) file. Entropy-based methods compute the variability of specific amino acid positions in an alignment of related protein sequences, allowing the identification of highly varying positions (Kalinina et al. 2004; Ye et al. 2006; Feenstra et al. 2007). As an example, SDPpred uses mutual information entropy scores to predict SDPs (Kalinina et al. 2004). Evolutionary-based methods use substitution matrices or phylogenetic trees to calculate residue-based variability scores (del Sol Mesa et al. 2003; Pazos et al. 2006; Capra & Singh 2008). Xdet (Pazos et al. 2006), a representative evolutionary-based method, calculates a similarity matrix for each alignment position by using a substitution matrix and, in parallel, computes functional similarities of protein sequences using GO terms or EC annotations. Finally, it assigns a score to each residue based on the Spearman-rank correlation between residue-based and functional similarity matrices. Xdet was recently improved to determine partner-specific SDPs (Pitarch et al. 2020). The feature-based methods perform feature extraction from each amino acid position by using sequence information. The constructed feature vectors are fed into a classifier such as random forest, support vector machine or neural network (Ahmad & Sarai 2005; Wong et al. 2015). As a feature-based method, Ahmad and Sarai proposed a Position Specific Scoring Matrix-based (PSSM-based) SDP prediction of DNA binding proteins (Ahmad & Sarai 2005). In this method, PSSMs were created for each protein sequence based on a reference dataset, where the rows represent residues of a target protein sequence and the columns correspond to 20 different amino acids. Here, each residue is represented as a feature vector using its and its neighbors’ conservation scores. Finally, created feature vectors are fed into a neural network classifier, to categorize the input residues as an SDP or non-SDP for DNA binding.

As the sequence-based SDP prediction methods do not use complex input data, they are computationally efficient. On the other hand, the methods proposed so far mostly trained with only small sequence datasets, using standard machine learning algorithms, challenging the accuracy of these methods.

### What could structural information add to SDP prediction?

According to core-rim interface model, protein-protein interfaces can be divided in two regions: the core, dominated by hydrophobic and conserved amino acids; and the rim, whose interactions involve solvent exposed and often non-conserved amino acids (Levy 2010). In a recent work of Ivanov *et al*., the core-rim interface definition was utilized to discriminate SDPs of four Paralogs protein families (Ivanov et al. 2017). Here, the authors structurally modeled and analyzed all possible interactions of these families, for which the experimental affinities were available. Their analysis showed that SDPs are found at the rim, where they make strong electrostatic (charge-charge) interactions (Chakrabarti & Janin 2002; Ivanov et al. 2017). Moreover, other groups utilized atomistic molecular dynamics (MD) simulations to trace the selective electrostatic inter-molecular interactions to pinpoint SDPs. As a prominent use of MD simulations to characterize SDPs, van Wijk *et al*. showed that a dynamic salt bridge network regulates specific ubiquitin-conjugating enzyme (E2s) and ubiquitin ligase (E3s) binding (van Wijk et al. 2012). Being at the rim of E2-E3 surface, the selective role of this salt bridge was validated by mutagenesis and yeast two-hybrid binding experiments. Another recent example highlights the grounds of the specific pairing of clustered protocadherins, which play an important role in neuronal development. In this work, Nicoludis *et al*. explored how molecular dynamics simulations and evolutionary couplings could be used together to dissect the dynamic factors, regulating specific protocadherin polymerization (Nicoludis et al. 2019).

Compared to the sequence-based SDP prediction methods, the above-listed structure-based approaches can provide a refined and experimentally testable SDP set. However, these structure-based methods require expertise in computational structural biology tools and, depending on the size of the system, they could be too computationally intensive.

### Which SDP prediction method: sequence- or structure-based?

As sequence- and structure-based methods have different advantages, we chose a model system to map the prediction landscape of these approaches. For this, we concentrated on a paralogous protein receptor tyrosine kinase family (TAM), consisting of Tyro3, Axl, and Mer proteins. TAM receptors, like the other receptor tyrosine kinases, are activated through their interactions with extracellular proteins, triggering receptor dimerization and autophosphorylation of their kinase domains (Rothlin & Lemke 2010). Earlier studies identified two proteins, the growth arrest-specific protein 6 (Gas6) and vitamin K-dependent protein S (Pros1) as TAM receptor agonists (Hafizi & Dahlbäck 2006). The binding of these ligands to TAM lead to downstream activation of diverse signaling pathways (Wium & Paccez 2018).

TAM receptor sequences are 52-57% similar, where their receptors are of 40% sequence similarity. Within family, Pros1 can bind to Tyro3 and Mer, while it cannot bind to Axl. Gas6, on the other hand, can bind to all three receptors with the highest affinity towards Axl (Hafizi & Dahlbäck 2006; Yanagihashi et al. 2017). Among the different combinations, the Axl:Gas6 interaction is particularly interesting given its involvement in numerous types of signaling pathways (e.g tumor-cell growth, metastasis, epithelial to mesenchymal transition, drug resistance etc.) (Zhu et al. 2019). Relatedly, Axl aberrant regulation was shown to lead to different types of cancer and infectious diseases (Van Der Meer et al. 2014), as well as to promote SARS-CoV-2 entry into cell (Wu et al. 2017; Wium & Paccez 2018; Wang et al. 2021). Although the structure and binding profile of Axl are both known, its specificity-determining positions remains unknown. As such, to help close this knowledge gap, we used three sequence-based SDP predictors, SDPred, Multi-RELIEF (feature-based), and Sequence Harmony (entropy-based) to map Axl SDPs. In addition, we carried out structure-based analyses of Axl:ligand interfaces, both using simple refinement calculations and extensive molecular dynamics simulations. Our work provides a key report on the comparative performance of SDP prediction approaches and paves the way for the development of therapeutic molecules capable of combatting Axl misregulation in different types of diseases.

## RESULTS

### A close look into the Axl:Gas6 interactions

TAM receptors share two immunoglobulin (Ig)-like, two fibronectin type III domains (FNIII), followed by a single-pass transmembrane helix, and an intracellular kinase domain (Figure 1A). TAM ligands, Gas6 and Pros1 contain an N-terminal gamma-carboxyglutamic acid (GLA) domain, four epidermal growth factor-like (EGF) repeats, and two laminin G (LG)-like domains (Figure 1A). The crystal structure of the Axl:Gas6 interaction is the only TAM:ligand structure at hand (PDB ID: 2C5D (Sasaki et al. 2006). This structure demonstrates that the two Ig-like domains of Axl interact with the two LG-like domains of Gas6, without involving any receptor-receptor or ligand-ligand contacts (Figure 1B). Axl and Gas6 interact with each other, through two symmetric copies of major and minor interfaces, which individually bury 2366 Å^2^ and 765 Å^2^ surface areas (Figure 1C). While the minor interface is highly conserved across the TAM subfamily, the major interface is not. The major interface is spatially segregated into a frontal site, involving a series of charged residues, and a hydrophobic distal site (Figure 1B, D) (Sasaki et al. 2006). The segregated characteristics of the major interface of Axl contributes to its ligand selection, as well as to its high affinity toward Gas6 interaction (Sasaki et al. 2006). Therefore, for studying the ligand selectivity of Axl, in this work, we focused on the major Axl:Gas6 interface (Figure 1B-inset).

**Figure 1:**
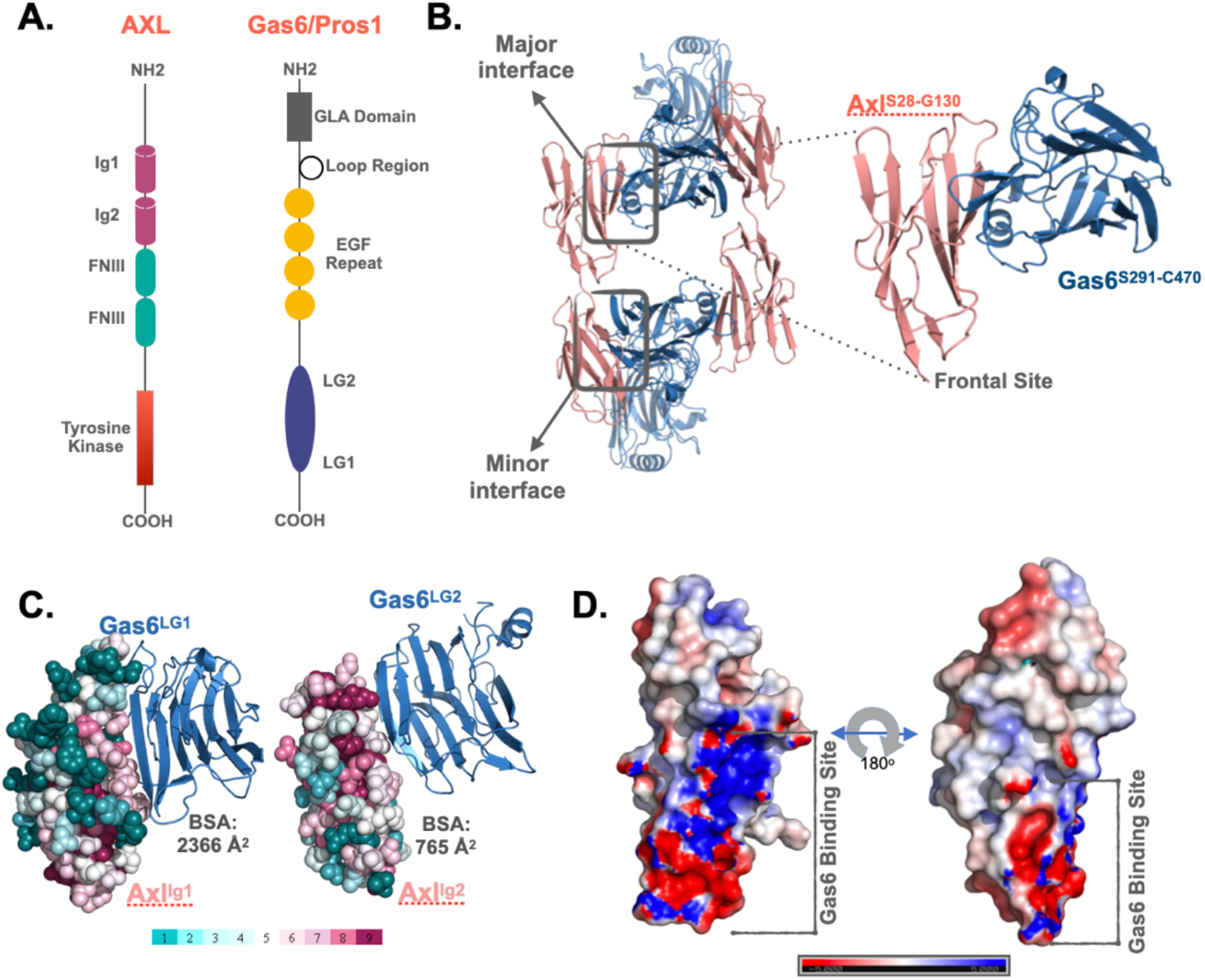
**A**. The domain organization of TAM family and its ligands, Gas6 and Pros1. TAM family consists of Ig1, Ig2, two FNIII, and tyrosine kinase domains (Lemke, 2013; Linger, Keating, Earp, & Graham, 2008). Gas6 and Pros1 are composed of GLA domain, loop region, EGF Repeat, and LG2, LG1 domains (Lemke, 2013; Linger et al., 2008). **B**. Axl(Ig1-Ig2):Gas6(LG1-LG2) interaction involves two interfaces: The major interface is formed between Axl-Ig1:Gas6-LG1 and the minor one is established among Axl-Ig2:Gas6-LG1 (PDB ID: 2C5D, (Sasaki, Knyazev, Clout, Cheburkin, Göhring, et al., 2006a)). The inset represents the charged frontal side of the major interface (Axl is depicted in pink cartoon, whereas Gas6 is represented in purple cartoon). **C**. Conservation scores of Axl residues predicted via ConSurf webserver (Glaser et al. 2003; Landau et al. 2005; Ashkenazy et al. 2016). **D**. Electrostatic potential of Axl:Gas6 interacting site. The color scale ranges from -5 (red) to 5 (blue). One side of Axl’s Gas6 binding surface is heavily charged, while the other side is composed of neutral amino acids.

### The sequence-based Axl SDP predictions agree in one amino acid position

Among the available sequence-based SDP predictors, we selected three to probe Axl ligand selectivity (Supplementary Table 1). These algorithms, i.e., SDPpred (http://bioinf.fbb.msu.ru/SDPpred/), Sequence Harmony (https://www.ibi.vu.nl/programs/seqharmwww/), and Multi-RELIEF (http://zeus.few.vu.nl/programs/multirelief/) were selected based on their widespread use and their availability as a web service (Kalinina et al. 2004; Feenstra et al. 2007; Ye et al. 2008). The mammalian (human, mouse, rat, pig, chimpanzee) TAM Ig1 sequences were retrieved from UniProtKB (The UniProt Consortium 2018). These sequences were then grouped into two, as Axl and Tyro3 & Mer. The MSA of each group was constructed with Clustal Omega (Sievers & Higgins 2017). For each approach, MSAs were formatted according to the requirements of the webservers. As earlier studies showed that SDPs tend to reside at the rim of a protein-protein interface, we filtered out the sequence-based SDP predictions by keeping the positions corresponding to the rim of the Axl:Gas6 complex (Ivanov et al. 2017). As a result, SDPpred predicted 19 SDPs, five of which (THR46, ARG48, GLN50, ASP84, LYS96) are at the rim of Axl:Gas6. The majority of the SDPpred predictions correspond to the non-interacting regions of the Axl:Gas6 complex (Duarte et al. 2012). The same trend was observed for Multi-RELIEF, which contained two rim Axl amino acids out of 15 SDP predictions (ARG48, GLU70). In the case of Sequence Harmony, the minority of the predictions (4/15) were found at the rim of Axl:Gas6 (THR46, ARG48, GLN50, LYS96). As such, the combined Axl SDP list, predicted by these three webservers is THR46, ARG48, GLN50, GLU70, ASP84, LYS96, where they all agree only on ARG48. The complete list of the SDPs predictions is provided under Supplementary Table 2.

**Table 1:** **A**. Positional comparison of GLU59, ARG48, GLU70, ASP73, GLU83. **B**. Comparison of GLU455^Gas6^, ARG414^Gas6^, ARG313^Gas6^, ARG467^Gas6^ residues with their Paralog counterpart.

| **A.** |  |  | **B.** |  |
| --- | --- | --- | --- | --- |
| **AXL** | **TYRO3** | **MER** | **GAS6** | **PROS1** |
| ARG48 | ASN63 | ASN114 | ARG310 | LYS314 |
| GLU59 | ASP73 | THR127 | ASP455 | ALA460 |
| GLU70 | GLN85 | LEU138 | ARG414 | ASN418 |
| ASP73 | ASP87 | HIS141 | ARG313 | LEU317 |
| GLU83 | - | ASP151 | ARG467 | ASN472 |

### Structural data revealed that Axl’s ligand selectivity is regulated by electrostatic interactions

To study Axl SDPs structurally, we modeled the three-dimensional structure of Ig1:LG1 Axl:Pros1 complex, to use it as a negative (non-binder) control. We refined the two Axl:ligand complexes with HADDOCK 2.2 webserver (https://alcazar.science.uu.nl/services/HADDOCK2.2/) (Van Zundert et al. 2016). When used for refinement, HADDOCK performs several independent short molecular dynamics simulations of the protein complex in explicit solvent. The best four refined Axl complexes, ranked by HADDOCK score differ mostly in interface electrostatics: Axl:Gas6 was ∼3.6 times higher than for Axl:Pros1 (-635.8±29 kcal/mol vs. -173.5±21 kcal/mol) (Figure 2A). Other interface features and energy terms, such as buried surface area and van der Waals energies, are comparable between the complexes. These results underscore the notion that Axl affinity and selectivity are mainly driven by the electrostatics interactions.

**Figure 2:**
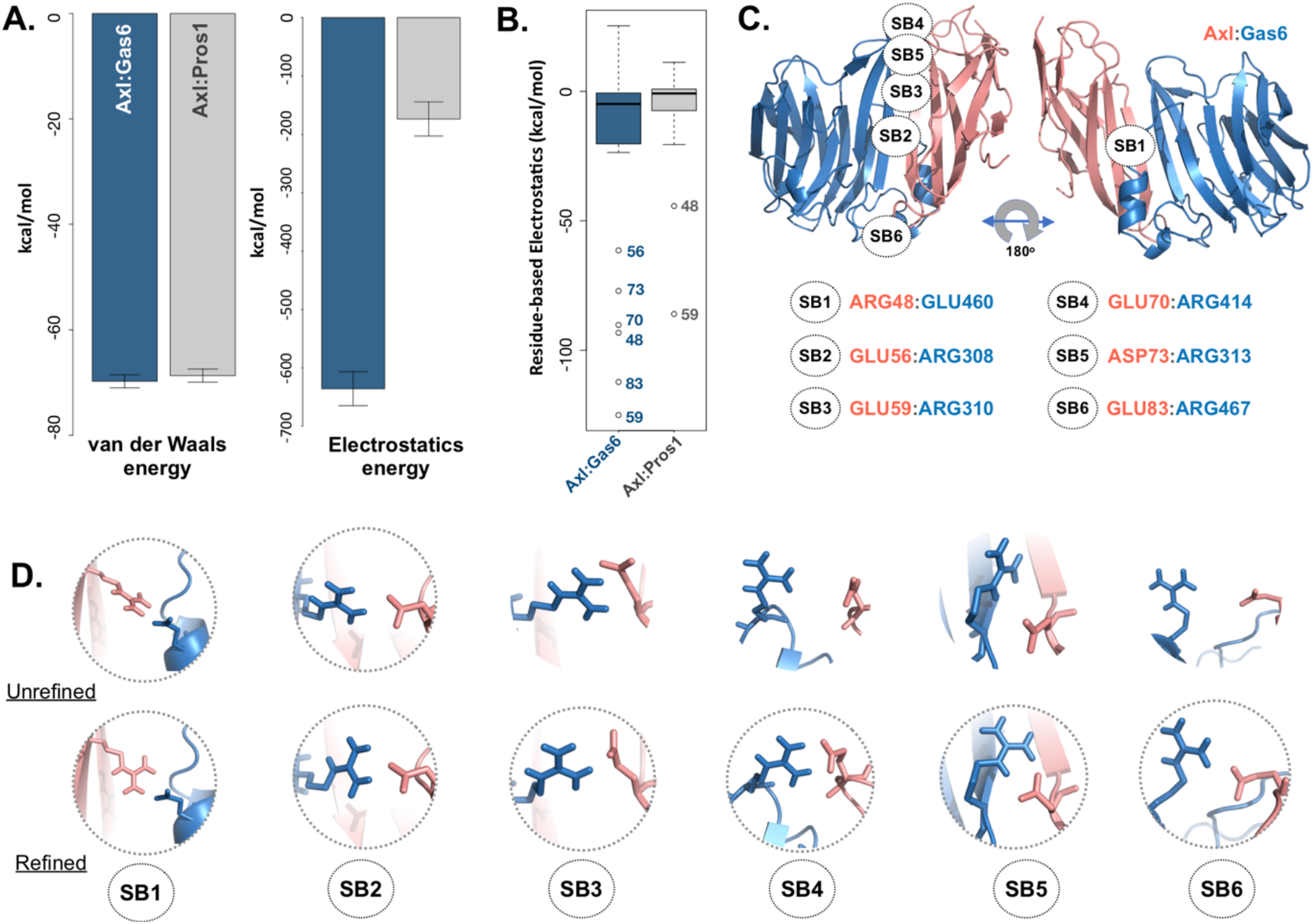
**A**. The mean van der Waals and electrostatics energetics of the top scoring Axl:Gas6 and Axl:Pros1 major interfaces calculated by HADDOCK. The mean van der Waals energies of Axl:Gas6 (blue) and Axl:Pros1 (gray) are -69.8±1.2 kcal/mol and -68.9±2.2 kcal/mol, respectively. The mean electrostatics of Axl:Gas6 and Axl:Pros1 are -635.8±29.3 kcal/mol and -173.5±21.4 kcal/mol, respectively. **B**. Residue-based electrostatics contribution of Gas6-facing Axl residues. Axl residues behaving differently than the rest of the population are ARG48, GLU59, GLU70, ASP73, GLU83 in the case of Axl:Gas6 (blue), and GLU48, GLU59 in the case of Axl:Pros1 (gray). The whiskers were computed with a whisker length of 2xIQR. **C**. The distribution of the salt bridges (SBs) across Axl:Gas6 interface. Axl is represented in light pink and Gas6 in marine blue. Only SB1 is located on the back and rather neutral side of the complex. This and all the structural images were generated with PyMOL molecular visualization software (Schrödinger, LLC. 2015). **D**. The arrangement of the potential SB-making residues in the case of crystal (first row, pdb id: 2C5D) and HADDOCK-refined (second row) Axl:Gas6 complex. The pairs forming SBs are encircled.

We also calculated per-residue electrostatics of interfacial Axl residues: 31 for the Axl:Gas6 complex and 27 for Axl:Pros1 (Figure 2B). In the case of Axl:Gas6, Axl ARG48, GLU56, GLU59, GLU70, GLU73, GLU83 contribute the most to the interface electrostatics (Figure 2B). These residues reside at the rim of the Axl:Gas6 complex, where they form six different salt bridges (Figure 2C, Supplementary Table 2). As laid out in earlier findings, the SDPs are rim amino acids, capable of making charge-charge interactions, which makes these salt bridge forming residues perfect SDP candidates. Among these six salt bridges (SBs), SB2-6 are located on the charged frontal side of the complex (Figure 2C, left). When we analyzed Axl:Pros1 interface, we found that SB1 and SB3 (mediated by ARG48, GLU59) were present there too. We therefore eliminated ARG48, GLU59 from the initial Axl SDP list, leaving GLU56, GLU70, ASP73, GLU83 Axl residues as the strongest SDP candidates. Here, we should note that SB3-SB6 were not present in the Axl:Gas6 crystal structure (Figure 2D). The proper establishment of these salt bridges could only be secured after HADDOCK refinement, highlighting the importance of dynamics information in the characterization of SDPs.

### Tracing the dynamic interaction profiles of Axl complexes suggests four salt-bridges as SDPs

To explore the time-dependent interaction profiles of Axl:Gas6 and Axl:Pros1, we carried out molecular dynamics (MD) simulations of the complexes (Van Der Spoel et al., 2005). For each complex, we ran four independent MD simulations, totaling 1.6 microseconds. The analysis of these trajectories showed that the Axl:Gas6 complex is more stable than Axl:Pros1, as reflected in the lower mean root mean square deviation (RMSD) of atomic coordinates to the average structure (0.15±0.01 nm vs. 0.23±0.03 nm) (Supplementary Figure 1), and also in the lower and more stable radius of gyration profiles (Supplementary Figure 2). To perform a more in-depth analysis of the interactions between Axl and its ligands, we calculated the number of inter-molecular hydrophobic, hydrogen bonds and salt bridges with the *interfacea* python package (https://github.com/JoaoRodrigues/interfacea/tree/master) (Figure 3A). When we pooled the interface data for each complex, we observed that Axl:Pros1 contains a fewer number of contacts in all interaction types. The most significant difference between Axl:Gas6 and Axl:Pros1 interaction distributions were observed for salt bridges. Axl:Gas6 trajectories reflected, on average, four to five stable salt bridges, where this number drops to two in the case of Axl:Pros1 (Figure 3A, right panel). We then looked for the salt bridges, which were seen in four different trajectories consistently for more than 25% of the simulation time (Supplementary Table 3). Here, our assumption was that the SDP positions should form stable salt bridges within a trajectory and should be observed consistently across four trajectories. These criteria left us with five salt bridges, four of which were the same as the ones selected by HADDOCK: GLU70:ARG414^Gas6^ (SB4), ASP73:ARG313^Gas6^ (SB5), GLU83:ARG467^Gas6^ (SB6), GLU59:ARG310^Gas6^ (SB3) (listed in the decreasing observation frequency, as in Figure 2B). GLU56-mediated SB2, coming from our HADDOCK analysis was observed only in one replica, indicating that it could be coincidental. As another surprising outcome, ARG48 of SB1 formed a stable and consistent salt bridge with ASP455^Gas6^, instead of GLU460^Gas6^, which was suggested by HADDOCK refinement. Interestingly, GLU460^Gas6^ has a GLU correspondence on Pros1, while alanine is found at the position of ASP455^Gas6^ in Pros1 (Table 1B). In the case of Axl:Pros1, only GLU59^Axl^:LYS314^Pros1^ was observed in a statistically significant manner, which corresponds to SB3 of Axl:Gas6 (Figure 3B, Supplementary Table 3). In the light of these results, we identify Axl SDPs as ARG48, GLU70, ASP73, GLU83 from these MD simulations.

**Figure 3:**
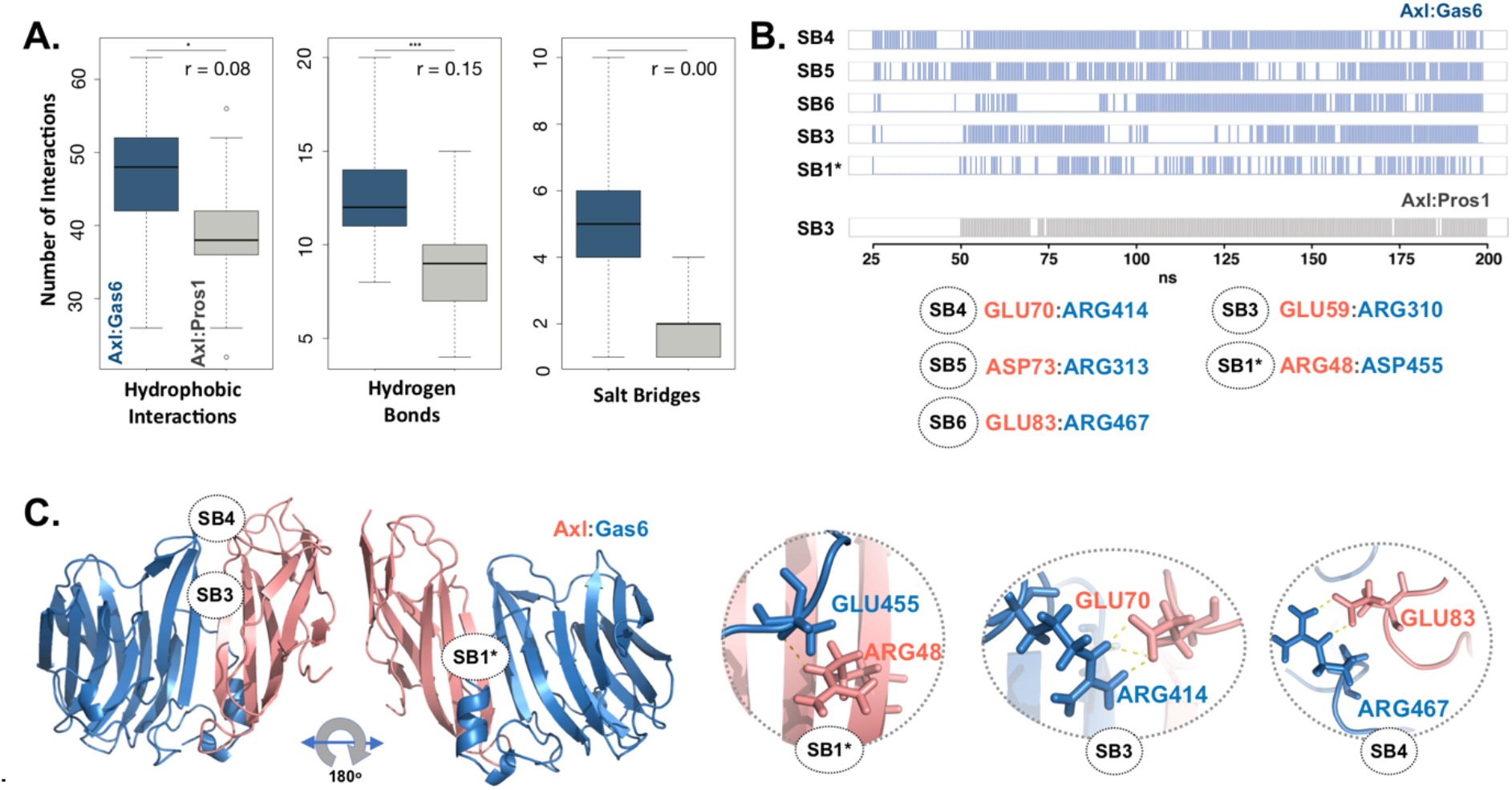
**A**. The distributions of hydrophobic contact (left), hydrogen bond (middle) and SB numbers (right) for Axl:Gas6 (marine blue) and Axl:Pros1 (light gray) simulations. The distributions are interpreted with box-and-whisker statistics. The associations between two Axl:Gas6 and Axl:Pros1 distributions were reflected in r values, according to which the SB distributions do not have anything in common. **B**. Consistently and stably observed SBs formed in Axl:Gas6 and Axl:Pros1 simulations. Each row indicates the observation frequency of the indicated SB. The frequency data came from replica 1 simulations (Supplementary Table 3). The SB numbering follows the ones used in Figure 2C. SB1 is denoted with * as in this case ARG48 uses a different partner than observed in Figure 2C. **C**. The positioning of potential SDPs on Axl:Gas6 structure predicted by molecular dynamics analysis.

### Evolutionary insights into the structure- and dynamics-based SDP predictions

If ARG48, GLU70, ASP73, GLU83 are Gas6-selective, their positions should be substituted to distinct amino acids in Tyro3 and Mer. The same should hold for Pros1 counterparts of GLU455^Gas6^, ARG414^Gas6^, ARG313^Gas6^, ARG467^Gas6^ (Figure 3B). To test this hypothesis, we performed an across-paralog sequence comparison (Table 1). The alignments between the ligand proteins showed that the charged residues in Gas6 involved in salt bridge formation with Axl were not conserved in Pros1. Besides, they were replaced with uncharged-polar or nonpolar residues, as in ^Gas6^ARG313LEU^Pros1, Gas6^ARG414ASN^Pros1, Gas6^ASP455ALA^Pros1, Gas6^ARG467ASN^Pros1^. On the other hand, ASP73^Axl^ was found to be conserved in Tyro3. We, therefore, eliminated ASP73 from the final Axl SDP list. Our across-ortholog comparison reveals that ARG48, GLU70, GLU83 are all positionally conserved, further supporting the SDP candidacy of these positions (Supplementary Figure 3). The distribution of the final list of salt bridges formed by these residues are illustrated in Figure 3C.

## DISCUSSION

In this work, we probed three sequence-based SDP predictors, SDPred, Multi-RELIEF, and Sequence Harmony to map a clinically interesting protein’s, Axl SDPs. Next to the sequence-based methods, we also carried out structure-based analyses, where we dissected Axl interactions with its ligands, which came out of simple refinement and extensive MD simulations. This research lays out a critical report on the comparative performance of SDP predictors.

As the primary outcome of this exercise, we found out that the sequence-based SDP predictors largely overpredict the potential SDP positions. Hence, we used available literature data to filter out the structurally non-viable SDP candidates. As a result, the three methodologies we used suggested six SDP positions combined, where they only agreed on ARG48. Our HADDOCK-based approach also suggested ARG48 as a strong electrostatic contributor to Axl:Gas6 interface. Though, here, we had to eliminate ARG48 from the potential SDP list, as it significantly contributed to the Axl:Pros1 interaction energetics too. Elaborate MD simulations were necessary to rescue ARG48’s SDP candidacy. We observed that during the MD simulations, ARG48 forms a new salt bridge, which was neither observed in the crystal, nor in the HADDOCK refined complexes. In the end, HADDOCK refinement proposed four selective SBs, three of which were supported by the MD simulations. Checking the evolutionary variance and conservation of MD-deduced SDP positions left us with ARG48, GLU70, and GLU83 as the strongest Axl SDP candidates. Strikingly, Multi-RELIEF (plus the rim information) could predict two of these (ARG48, GLU70) without running any extensive calculations.

### How can we validate our Axl SDP predictions?

We aimed at finding an orthogonal approach to validate the Axl SDPs. For this, experimentally, one could mutate the predicted SB-forming Gas6 residues to Pros1 counterparts or vice versa, followed by the measurement of receptor:ligand binding affinity changes. This would be the utmost way to unambiguously validate the Axl SDP predictions. However, accessing such specific experiments would not be within the reach of many computational biologists. Therefore, to probe the significance of Gas6-selective Axl residues, here, we artificially mutated four selectively SB forming Gas6 residues their Pros1 counterparts, and vice versa, by using EvoEF1 (https://github.com/tommyhuangthu/EvoEF) (Pearce et al. 2019) (Figure 4). EvoEF1 is a machine learning approach, poised to calculate the impact of point mutations across protein-protein interfaces. According to EvoEF1, Axl:Gas6 interaction stability was significantly reduced when individual and combined Gas6-to-Pros1 and Gas6-to-alanine mutations were imposed (Figure 4). On the other hand, individual and combined Pros1-to-Gas6 mutations led to a significant increase in the stability of Axl:Pros1 complex. These findings underscores the vitality of the selective Gas6 positions to the formation of Axl:Gas6 complex.

**Figure 4:**
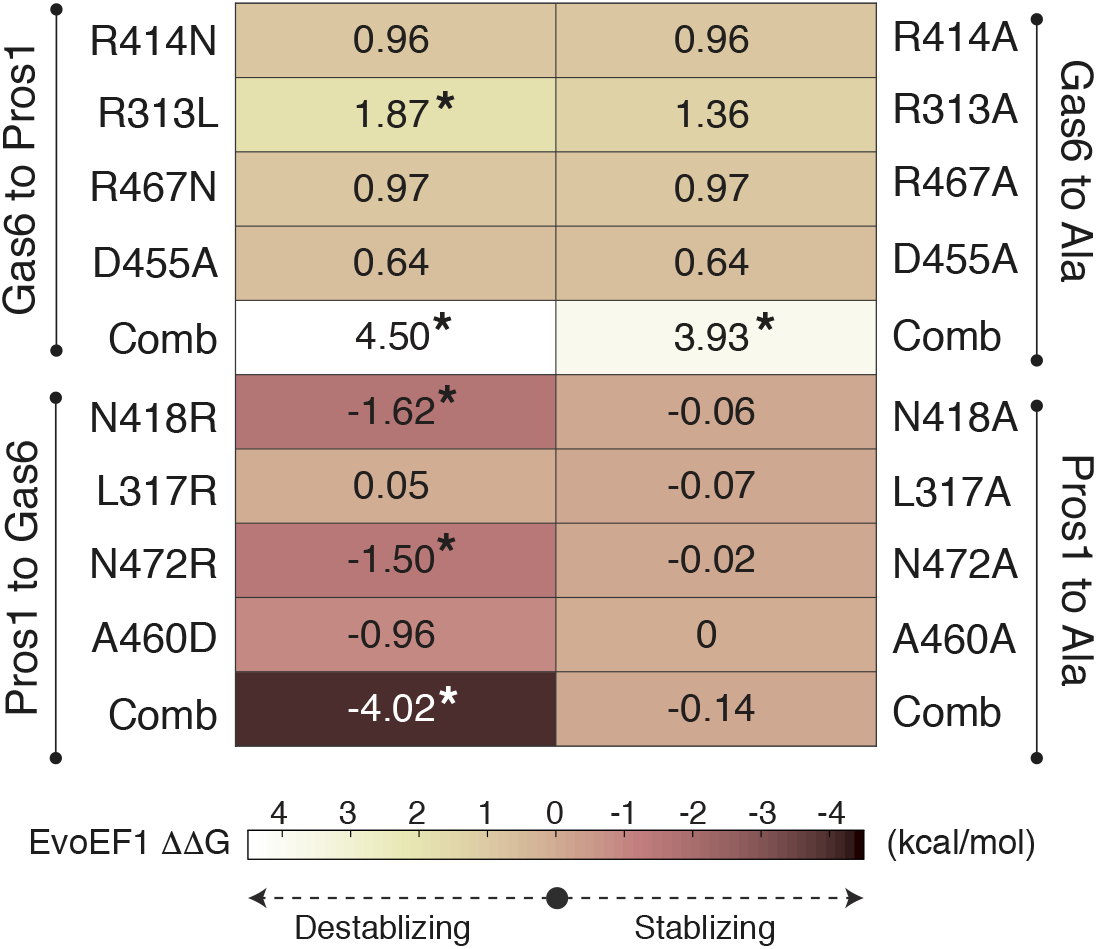
EvoEF1 ΔΔG predictions for the selective Gas6-to-Pros1 or Pros1-to-Gas6 mutations. ARG414, ARG313, ARG467, and ASP455 correspond to SB4, SB5, SB6, and SB1* forming residues, as presented in Figure 3B. The significant changes are marked with *. The stabilizing and destabilizing mutation color scheme ranges from -4 (brown) to 4 (yellow). Gas6-to-Ala and Pros1-to-Ala mutations were run as control.

### What awaits the SDP prediction field in the future?

Given the importance of specificity determining positions for clinical purposes, it is essential to use an economically feasible and accurate predictor. To this end, using machine learning (ML) methodologies in SDP prediction is very suitable, as ML tools would allow calculating dozens of SDP predictions in seconds. From the accuracy point of view of though, current ML-based approaches face many challenges. As an example, the majority of the sequence-based SDP prediction methods require a precalculated MSA file, together with subfamilies or subgroups definition. Here, caution should be taken as different MSA algorithms produce different alignment results based on varying parameters, which in the end will affect the final SDP list. Besides, dividing protein families into subfamilies requires expert knowledge. As another important limitation, the experimentally determined SDP datasets are rather limited, which, in turn, prevents creating large-scale training of the feature-based methods. Construction of such large-scale SDP training datasets will make it possible to use deep learning algorithms, which have outperformed state-of-the-art methods in similar problems (LeCun et al. 2015). The structure-based methods, as presented in this work under the umbrella of HADDOCK and MD simulations, could offer a refined SDP list, which can be tested experimentally. These approaches, however, take much longer time as they explicitly use structures and calculate forces acting on these structures. As an example, HADDOCK refinement of complexes can take up to half an hour, depending on the available computing resources. MD simulations, on the other hand, can take up to days or weeks, based on the dedicated number of computing cores used. Considering the pros and cons of both methodologies, it is evident that new and novel SDP prediction methods, which combine the advantages of both sequence- and structure-based method, should be developed. However, until then, we propose the structure-based identification of SDPs by HADDOCK, in combination with Multi-RELIEF, which could deliver the MD-determined SDP positions without the requirement of heavy calculations.

## METHODS

### Sequence-based Methods

**SDPpred** is an entropy-based SDP prediction method which utilizes mutual information to determine well-conserved residues (i.e., SDPs) within the same groups but differ between them. The equation to mutual information score for a column p in the alignment is given below:

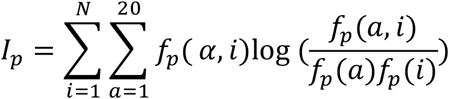

In this equation, *N* is the number of specificity groups, *a* is the amino acid type, *f*_*p*_(*i*) ratio of protein sequences belonging to group *i. f*_*p*_(*a*)is the number of occurrences of residue *a* in the whole alignment at position *p. f*_*p*_(*a, i*) is the number of occurrences of residue *a* in group *i*at position *p*. SDPpred calculates column-wised scores for each position in the MSA and outputs SDPs over the protein sequences.

**Multi-RELIEF** a machine-learning based SDP prediction method which employs RELIEF algorithm to identify specificity determining residues (Ye et al. 2008) (Kononenko 2005). Multi-RELIEF algorithm requires predefined groups and MSA as input. At each iteration, a random sequence is selected, and the weight of each residue is updated using the RELIEF algorithm by considering the most similar sequence from the same group and the most similar sequence from the other group. The algorithm outputs a weight vector whose length is the same as the number of positions in the alignment and higher weights indicates the higher probability of being SDP for the corresponding position.

**Sequence Harmony** is another entropy-based SDP prediction method (Feenstra et al. 2007). It takes MSA and two user-specified groups as input and calculates relative entropy scores for each residue that shows degree of conservations. Sequence Harmony provides ranking of the entropy scores as outputs. Sequence Harmony and Multi-RELIEF methods are merged under the Multi-Harmony web server at https://www.ibi.vu.nl/programs/shmrwww/ (Brandt et al. 2010).

### Template-based modeling of Axl:ligand complexes

LG1 domain of Pros1 was modeled with i-TASSER (Roy, Kucukural, & Zhang, 2010). Pros1-to-Gas6 structural alignment was carried out with FATCAT web-tool (Y. Ye & Godzik, 2003) (by using the Gas6 coordinates of 2C5D). The final Axl:Pros1 coordinates were visualized and saved in PyMOL (Schrödinger, LLC. 2015). All Axl:ligand complexes were water refined with HADDOCK2.2 web server (van Zundert et al., 2016). The standard HADDOCK refinement protocol samples 20 models. These models slightly differ from each other as each one is refined with a molecular dynamics simulation starting from a different initial velocity. In the end, the generated models are ranked with the HADDOCK score, which is a sum of electrostatics (elec), van der Waals (vdW) and desolvation terms (desolv): 1.0. vdW+ 0.2. elec + 1.0. desolv. The top ranking four models, i.e. the best four models with the lowest HADDOCK scores, are offered as the final complex states. We generated 200 refined structures for each Axl:ligand complex. The top four ranking models were isolated as the final solutions.

## HADDOCK-based energy analysis

HADDOCK reports van der Waals (vdW) energy, electrostatics energy, desolvation, total HADDOCK Score, as well as BSA of each models. HADDOCK also calculates residue-based energy scores of each complex (expressed in electrostatics, vdW and electrostatics+vdW), deposited in *ene-residue*.*disp* file (can be found under HADDOCK output folder: structures/it1/water/analysis). This file describes electrostatics, vdW and electrostatics+vdW contributions of each interface amino acid to the formation of the intermolecular interaction. Here, we considered electrostatics scores of Axl interface amino acids. To generate HADDOCK-related plots, “lattice” library and levelplot function of R were used (https://www.rdocumentation.org/packages/lattice/versions/0.10-10/topics/levelplot).

## Molecular Dynamics Simulations

GROMACS 5.1.4 software and its tools were used to run molecular dynamics simulations (MD) and quality controls (e.g temperature, pressure, RMSD, Rg analyses) (Van Der Spoel et al. 2005). The AMBER99SB-ILDN force field (Lindorff-Larsen et al. 2010) was used to parameterize the protein molecules, while the TIP3P water model was used to represent the solvent (Jorgensen et al. 1983). The simulation was run in a rhombic dodecahedron unit cell. The minimum periodic distance to the simulation box was set to be 1.4 nm. The mdp simulation files were adapted from https://github.com/haddocking/molmod-data.

Before the production run, each complex was minimized in vacuum by using the steepest descent algorithm (Mandic 2004). They were then solvated with the TIP3P water, together with neutralizing ions (51 NA+ and 48 CL-ions were added to neutralize Axl:Gas6 while 58 NA+ and 49 CL-ions were added to neutralize Axl:Pros1). The relevant topology files were edited according to the newly included NA+ and CL-ions. The second cycle of energy minimization was performed on the solvated systems. The solvent and hydrogen atoms were relaxed with a 20 ps long molecular dynamics simulation under constant volume where the temperature is equilibrated to 300 K (NVT). This was followed by 20 ps long molecular dynamics simulation under constant pressure where the pressure is equilibrated to 1 bar (NPT). As a last step before the production run, position restraints were released upon reduction of its force constant from 1000 to 100, 100 to 10, and 10 to 0. To generate a parallel run of a given complex, random seed was changed before running the NVT step. The coordinates were written in every 10 ps. The integration time step was set to 2 fs.

For each complex simulation, upon reaching 200 ns, the periodic boundary conditions were corrected. The system was stripped off solvent and ion atoms. The Root Mean Square Deviations (RMSDs) were calculated by using the average coordinates as a reference. After leaving the equilibration periods out (25 ns for Axl:Gas6 and 50 ns for Axl:Pros1), 350 snapshots for Axl:Gas6 and 300 snapshots from Axl:Pros1 were extracted.

## Interface Analysis

The interfacial hydrophobic contacts, hydrogen and salt bridges were calculated with *interfacea* python library (https://github.com/JoaoRodrigues/interfacea) (Rodrigues 2019). *interfacea* classifies an inter-monomer interaction as hydrophobic, if there are at least two non-polar atoms within 4.4 Å. It considers a pairwise contact as a hydrogen bond, if a hydrogen donor (D) and acceptor (A) gets are within 2.5 Å. It then filters the matches for D-H-A triplets with a minimum angle threshold (default 120 degrees). Finally, it classifies an interaction as a salt bridge if there are oppositely charged groups within 4.0 Å. For the interface classification as core and rim, EPPIC webserver was used (Duarte et al. 2012). Core residues are the ones that are buried at least 95% in protein structure. The rest of interface residues are counted as rim residues.

For comparing different simulations, the box-and-whisker statistics were generated with the standard *boxplot* function of R. The salt bridges were classified as *stable* if they were observed for >25% of a simulation time. They were classified as *consistent* if they were observed to be stable in all simulations.

## Author Contributions

All authors contributed to the conceptualization of this work, as well as to the writing of the manuscript. Specifically: ASR and TK performed the sequence-based analysis. TK performed structure-based analysis. JPGLMR devised and developed the *interfacea* package. EK participated at all levels and supervised the work.

## Funding

This work was supported by the EMBO Installation Grant granted to Ezgi Karaca (project # 4421).

### Acknowledgements

We wholeheartedly thank Prof. Dr. Mehmet Öztürk and Drs. Tuğçe Batur for their assistance on Axl biology.

## Supplementary Material

Supplementary Material is submitted separately.

## Data Availability

All the relevant sequence- and structure-based input/output files are deposited at https://github.com/CSB-KaracaLab/Paralog_SDP

