## Supplementary Information for "Predicting the Specificity-Determining Positions of Paralogous Complexes"

SUPPLEMENTARY FIGURES

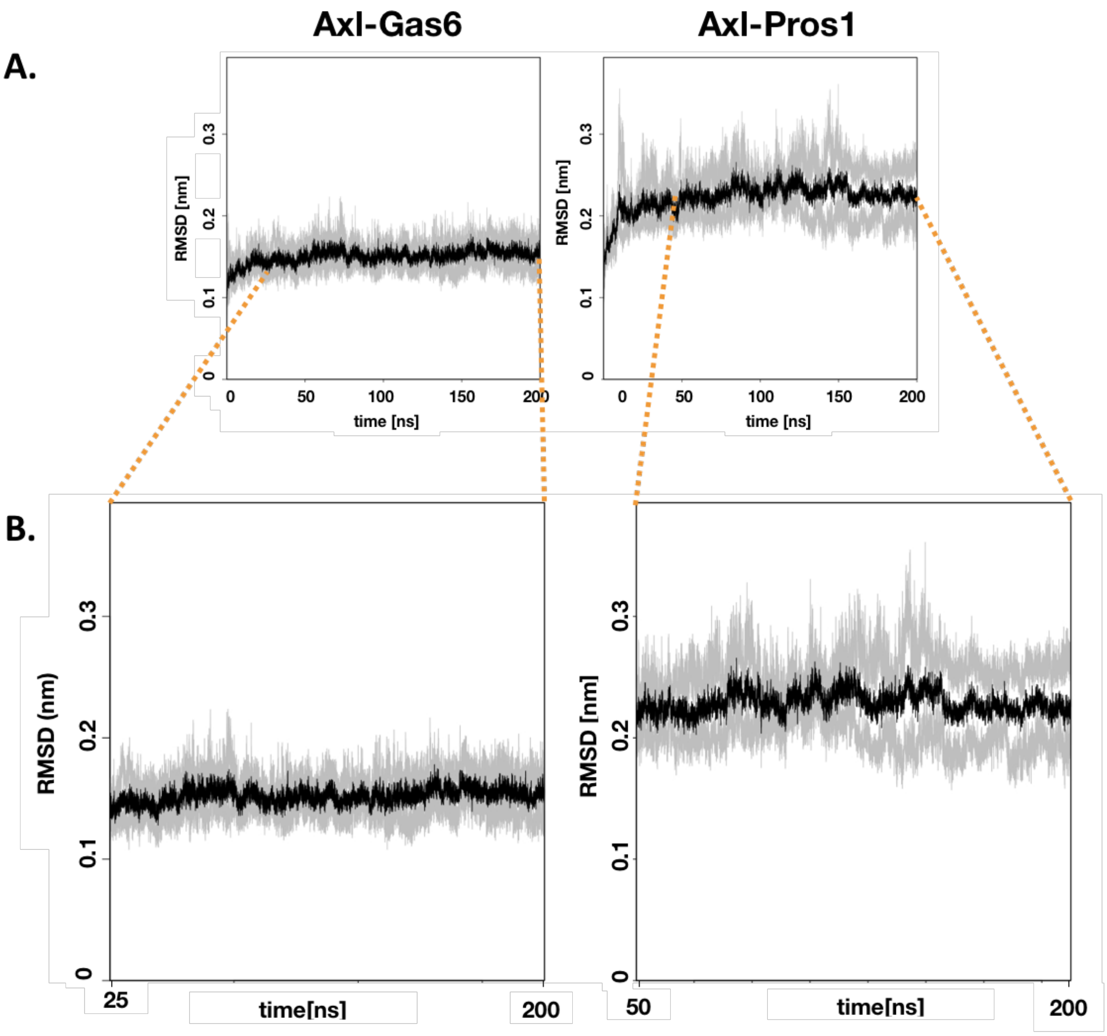


**Supplementary Figure 1:** Root mean square deviations (RMSDs) of Axl:Gas6 and Axl:Pros1 simulations from the average Axl:ligand structure. **A.** Axl:Gas6 simulations reach equilibrium in 25 ns, whereas Axl:Pros1 simulations reach it in 50 ns. **B.** During the production runs, Axl:Gas6 coordinates reflect 40% smaller mean RMSD (~0.15 nm) than Axl:Pros1 ones (~0.25 nm). Moreover, the RMSD values of Axl:Pros1 fluctuate between higher RMSD values (minimum: 0.15 nm and maximum: 0.36 nm for Axl:Pros1; minimum: 0.13 nm and maximum: 0.22 nm for Axl:Gas6).

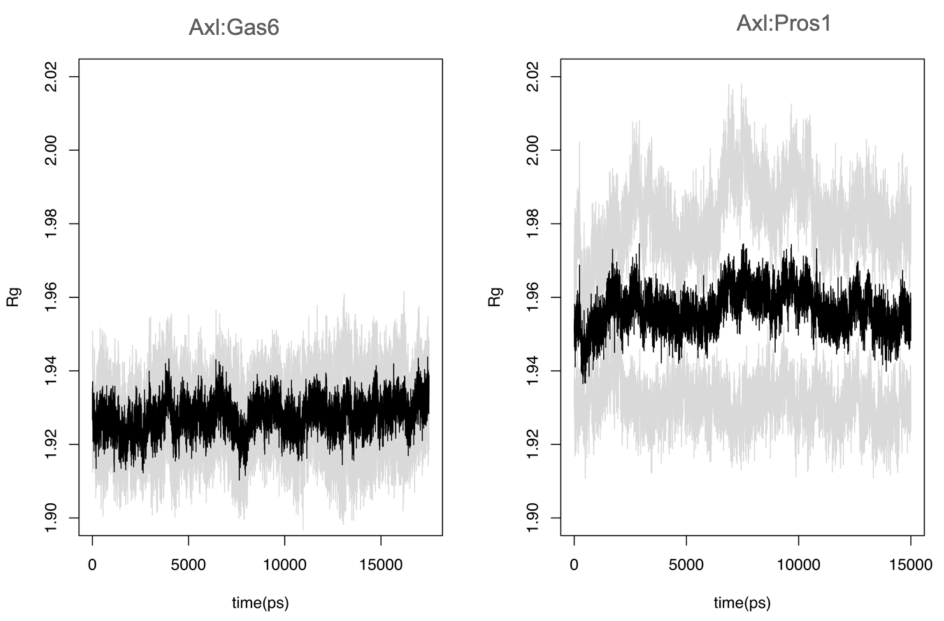


**Supplementary Figure 2: A.** The radius of gyration fluctuations (calculated over the backbone, expressed in nm) of Axl:Gas6 (left) and Axl:Pros1 (right) simulations.

**A.**

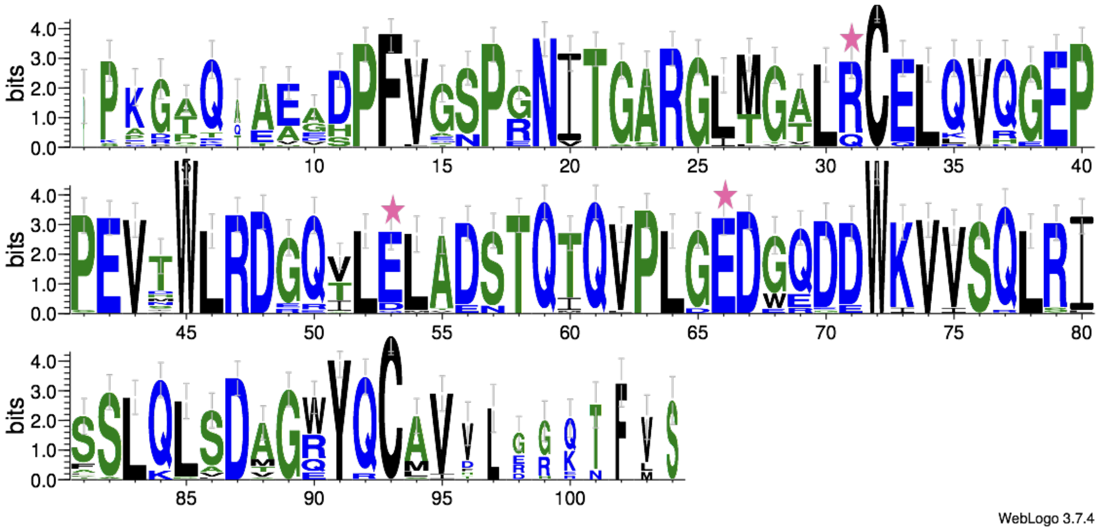

**B.**

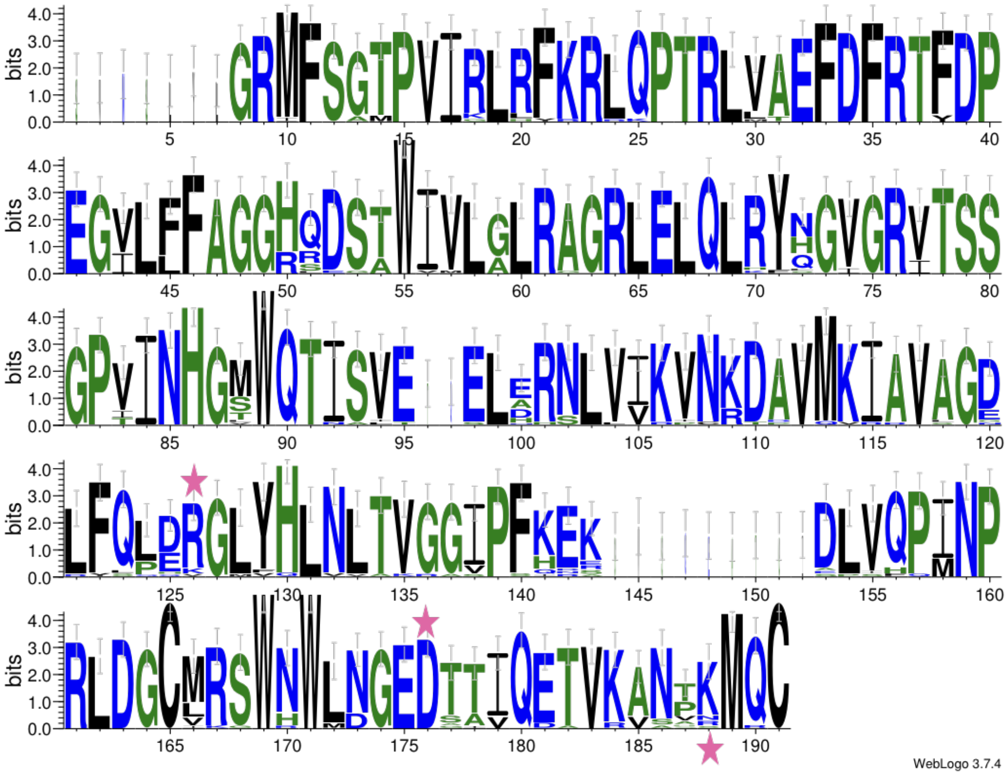

**Supplementary Figure 3:** Conservation of Axl SDPs and Gas6 residues across Axl and Gas6 orthologs

**SUPPLEMENTARY TABLES**

**Supplementary Table 3:** SBs formed in parallel **A.** Axl:Gas6 and **B.** Axl:Pros1 simulations. Each row indicates the observation frequency of the denoted salt bridge. The SB numbering follows the ones used in Figure 2C. The consistent and stable SBs are marked in bold.

| **A.** | **Axl** | **Gas6** | **Axl-Gas6-replica #1 (%)** | **Axl-Gas6-replica #2 (%)** | **Axl-Gas6-replica #3 (%)** | **Axl-Gas6-replica #4 (%)** |
|---|---|---|---|---|---|---|
| **SB4** | **70** | **414** | **81.14** | **34.57** | **55.41** | **70.57** |
| **SB5** | **73** | **313** | **72.86** | **81.71** | **69.43** | **76.86** |
| **SB6** | **83** | **467** | **60.29** | **48** | **68.57** | **71.14** |
| **SB3** | **59** | **310** | **59.43** | **32.86** | **83.14** | **56.86** |
| **SB1*** | **48** | **455** | **43.71** | **44** | **53.43** | **30.57** |
| SB7 | 70 | 312 | 30.57 | 40 | --- | --- |
| SB8 | 88 | 467 | 28.29 | 28 | 26 | --- |
| SB9 | 83 | 299 | --- | 33.42 | 60.29 | 34.86 |
| SB10 | 85 | 308 | --- | --- | 36.86 | --- |
| SB11 | 96 | 454 | --- | 44.71 | 27.71 | --- |
| SB2 | 56 | 308 | --- | --- | --- | 25.14 |
| SB13 | 48 | 460 | --- | 56.86 | --- | --- |
| SB14 | 59 | 414 | --- | 56.29 | --- | --- |

| **B.** | **Axl** | **Pros1** | **Axl-Pros1-replica #1 (%)** | **Axl-Pros1-replica #2 (%)** | **Axl-Pros1-replica #3 (%)** | **Axl-Pros1-replica #4 (%)** |
|---|---|---|---|---|---|---|

| | | | | | | |
|---|---|---|---|---|---|---|
| **SB3** | **59** | **314** | **97.67** | **97.67** | **94.33** | **94.33** |
| SB2 | 59 | 316 | -- | -- | -- | 66.66 |
| SB1 | 48 | 465 | 64.67 | 51.67 | -- | 33 |
